# An in-depth dataset of northwestern European arthropod life histories and ecological traits

**DOI:** 10.1101/2024.12.17.628846

**Authors:** Garben Logghe, Femke Batsleer, Dirk Maes, Tristan Permentier, Matty P. Berg, Dimitri Brosens, Stijn Cooleman, Pallieter De Smedt, Jonas Hagge, Jorg Lambrechts, Marc Pollet, Fons Verheyde, Dries Bonte

## Abstract

In response to the ongoing biodiversity crisis among arthropods, it is essential to implement efficient conservation strategies to safeguard both species diversity and the vital ecosystem services they provide. Developing such strategies requires reliable predictive models that can identify the species that are the most vulnerable to current and future threats, including those posed by climate and land-use change. Species life histories are central to these models, as they influence both population dynamics and spread rates. To support this effort, we compiled a dataset with key traits for arthropods based on several literature sources and expert knowledge. The dataset contains data on body size, life history, thermal niche and ecology for 4874 northwestern European species across 10 different orders. By gathering these essential trait data, we aim to create a robust foundation for predicting species vulnerability and anticipating shifts in arthropod communities in response to global change.

## Introduction

Arthropods are an exceptionally species-rich taxon that provide essential ecosystem services such as pollination or nutrient cycling (Stork, 2018; Yang & Gratton, 2014). Despite their critical importance, arthropods remain noticeably understudied in conservation research compared to vertebrates (Clark & May, 2002). This oversight is particularly concerning as these vital organisms are currently under threat from a widespread crisis referred to as “a death by a thousand cuts” (Wagner et al., 2021). The problem stems from a combination of global threats, including climate extremes, pollution, eutrophication, invasive species and urbanization, which have collectively led to significant declines in both the abundance and species richness of arthropods (Harvey et al., 2023; Wagner, 2020). The situation is expected to worsen with impending global change, putting arthropods at even greater risk (Hallmann et al., 2017; Seibold et al., 2019; Soroye et al., 2020).

A key strategy to increase arthropod community resilience against global change involves maintaining biotic complexity (Samways et al., 2020). This can be achieved locally by conserving both at-risk habitats and microhabitat heterogeneity (Toonen et al., 2022). On a regional scale, it requires restoring habitat quality and connectivity, enabling species with restricted environmental tolerances to track their optimal niches under global change (Travis et al., 2013). However, predicting which species will be most vulnerable and where conservation measures should be prioritized remains challenging. Existing predictive models are mostly based on correlative species distribution projections but neglect biological mechanisms such as demography, dispersal and interactions between species. Including these mechanisms is crucial for producing accurate forecasts, as merely extrapolating correlations between species’ range and climate is insufficient (Urban et al., 2016).

Implementing these biological mechanisms in predictive models requires detailed information on species’ demographic and ecological traits (Comte et al., 2024; Urban et al., 2016). Compiling such trait data for arthropods is challenging for two main reasons: many arthropod groups are less studied compared to vertebrates (Leather, 2013) or plants (Kleyer et al., 2008), and their high species richness, small sizes and hidden life styles make it difficult to quantify key traits. Furthermore, available trait data are mostly scattered across literature, with non-standardized measurements that hinder cross-taxon comparisons (Moretti et al., 2017).

To address this gap, we aimed to compile relevant trait data needed for predictive demographic models of arthropod diversity under global change pressures. We synthesised data on life history traits such as fecundity, dispersal ability and development time from various literature sources and expert judgement. Additionally, we included information on readily available ecological traits like habitat, feeding guild and thermal niche. With the exception of genetic adaption, our goal is to provide data on all necessary ecological components to construct reliable predictive models for forecasting the impact of global change on arthropod species (Urban et al., 2016).

## General description

### Purpose

This dataset summarizes life history and ecological traits for 4874 European arthropod species, aiming to construct a comprehensive dataset that could be incorporated in predictive models. The selection of traits was guided by four of the six mechanisms identified by Urban et al. (2016) that determine biological responses to climate change: dispersal, demography, species interactions and physiological processes. We are convinced that most of these mechanisms can also be integrated into models assessing the effects of other threats, such as land-use change. The first mechanism is dispersal, defined as the movement of a species from its place of birth to their location of reproduction (Clobert et al., 2009). The dispersal ability of a species is crucial for its ability to track suitable niches in new environments, making it a key component of this dataset (Travis et al., 2013). A second important mechanism is demography, determining population growth rates (Altermatt, 2010; Bommarco, 2001). It is here represented by the following life history traits: fecundity, development time, voltinism and lifespan. Overwintering stage was also included due to its strong connection with climate change-induced phenological shifts, which can significantly alter population dynamics (Marshall et al., 2020). A third mechanism is species interactions, acknowledging that species do not exist in isolation. Although it was not feasible to include direct measures of competition (e.g. functional responses) in this dataset, we did incorporate feeding guild and trophic range, which can be relevant in the context of global change (Burghardt & Tallamy, 2013; Mitchell & Litt, 2016). The fourth type of mechanisms is physiological processes, which are crucial to understand population change under climate change. To address this, we included realised thermal niches to indicate a species’ vulnerability to shifts in temperature regimes. Additionally, diurnality was included, as it may influence species’ responses to climate change (Levy et al., 2019). The final two mechanisms identified by Urban et al. (2016) – environment and evolution – were not included in this dataset due to practical constraints. Environmental factors are largely species-independent and should be sourced from other datasets. Nevertheless, we included habitat information, as it is essential for assessing the responses of specific communities. The evolution mechanism, which involves genetic variation, was omitted due to the lack of available data for most arthropod species. In addition to traits related to these six mechanisms, we included data on body size, overwintering stage and diurnality. Body size is often considered a “master trait” because it correlates with many other traits related to several of the key mechanisms (White et al., 2007; Woodward et al., 2005). Body size can determine maximal population densities (Currie, 1993; White et al., 2007), movement capacities (Hillaert et al., 2018; Logghe et al., 2024), fecundity (Berger et al., 2008) and thermal limits (Klockmann et al., 2017). The inclusion of body size in our data consequently allows it to be used as a correlated trait and proxy for potential missing data on specific mechanisms. With this selection of traits, we aim to facilitate the implementation of biological mechanisms in predictive models focused on arthropods, thereby enhancing our ability to forecast the impact of global change on these animals.

## Sampling methods

### Sampling description

Initially, we conducted an extensive search for literature sources that describe life history and/or ecological traits of arthropods. Our goal was to include as many arthropod orders as possible. However, we soon realized that to obtain sufficient data per taxon, we needed to focus on 10 specific orders (see Taxonomic coverage). The data were compiled in separate Excel sheets for each order. Each source was recorded on a separate line in the dataset, regardless of whether the species had already been included. This approach allowed us to consider multiple sources when estimating trait values for different species. Later, this information was merged into a single dataset (see below). We consulted 83 different literature sources (see Supplement 1 for a full list), including scientific papers, books and websites (Figure 1). Initially, the search for data was focused on existing datasets or books with extensive descriptions of specific groups, aiming to find sources with the most comprehensive information. Subsequently, we targeted specific papers on individual species or small groups to fill gaps in the dataset.

**Figure 1:**
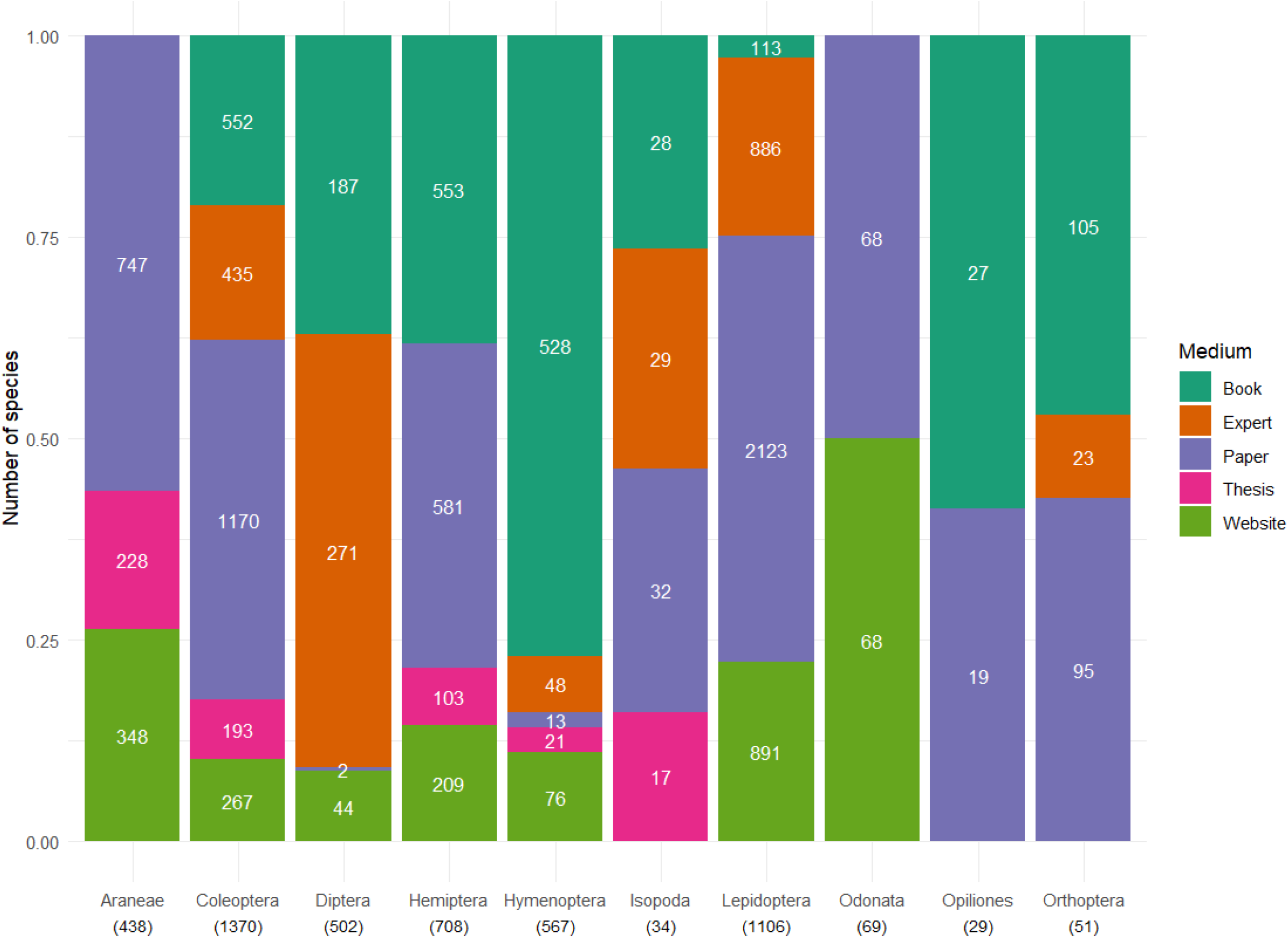
Overview of the proportional contribution of various data sources to each order. Absolute species counts are depicted by the white numbers within the graph. Total number of species per order are below order names between brackets.

In the next step, several authors contributed personal data, measurements and expert judgements on specific arthropod groups to address noticeable gaps in the dataset. These data points were added as separate rows to the raw datasets, with the contributing author’s name cited as the source. One of the most significant contributions from these experts was the estimation of dispersal ability, defined as a species’ potential to move from its birthplace to new areas. This estimation was invaluable, as dispersal data is often difficult to obtain from literature or simply unmeasured.

Additionally, we estimated the realised thermal niche for each species in the datasets. This was accomplished by overlaying species distribution data derived from GBIF (GBIF.org, 2023) with WorldClim climate data (Fick & Hijmans, 2017) in QGIS (QGIS Development Team, 2023). First, an R script was developed to automate the retrieval and processing of species occurrence from GBIF. The script utilizes the “rgbif” package (Chamberlain et al., 2024) to interface directly with the GBIF API, allowing for large-scale download of biodiversity data for multiple species. Specifically, the script matches a list of species names to their corresponding “speciesKey” in the GBIF taxonomic backbone. Once matched, the script initiates a data download using the “occ_download” function, filtering the data to include only records with geographic coordinates and excluding records with geospatial issues, null observations, preserved specimens, and fossils. After submitting the download request, GBIF provides a unique download key, which is used by the script to retrieve the dataset once it is available using the “occ_download_get” function. The data is imported into an R dataframe, where potential issues with name matching or data availability can be checked. The script then creates individual CSV files for each species, containing species names and geographic coordinates. For the next step, a Python (Van Rossum & Drake Jr., 1995) script was developed to run in QGIS. After loading a raster with annual mean temperatures (BIO1) derived from WorldClim, the script processes each CSV file to derive distribution data. The script creates a buffer around each species observation point (0.2 degrees) to expand it to a small area, ensuring that temperature data is averaged over a meaningful geographic range rather than at a single point. The zonal statistics tool is then used to calculate temperature statistics (minimum, maximum, and mean) within each buffer area by extracting raster values (annual mean temperature) that overlap with the buffers. The resulting data is stored in a CSV file containing the three temperature measures, with one row per assessed species. Finally, temperature range was calculated by subtracting the minimum temperature from the maximum.

In the final step, we aimed to create a centralized dataset that consolidated all species traits into a single, averaged value per trait for each species. This involved merging the raw datasets with the data on temperature niches and then summarizing each trait per species using R. For continuous traits, such as body size, we averaged all available values from different sources into a single value per species. When sources provided separate values for males and females, we first averaged sex-specific values and then combined them with values from other sources to derive the final species value. For categorical variables, we adopted the value representing the most inclusive or highest category. For example, if a species was described as both omnivorous and carnivorous by different sources, we classified it as omnivorous in the final dataset. Similarly, if one source indicated that a species overwinters as larva and another source did not, we classified the species as overwintering in the larval stage in the final dataset. Averaging dispersal ability was particularly challenging due to the significant variation in dispersal modes and studied proxies (from morphology to behaviour). For example, dispersal ability in spiders is often experimentally assessed by ballooning propensity, whereas in beetles, wing load is the most prevalent measure. Different proxies of dispersal, and metrics of dispersal ability make it further impossible to directly average these values across and within taxa. For example, one source might use a scale of 1-9 for estimating the dispersal ability of a butterfly, while another might use a scale of 1-10. To address this, we rescaled the dispersal ability values to a relative scale between 0.1 and 1 within each order. For example, if a butterfly received the highest dispersal score from a particular source (e.g. 9 on a 1-9 scale), it was assigned a value of 1. Conversely, the species with the lowest dispersal score (e.g. 1 on a 1-9 scale) would receive a value of 0.1. This process was applied separately to each order, ensuring that every group included species with dispersal values ranging from 0.1 to 1. By rescaling and averaging dispersal ability within each order, we aimed to create relative dispersal values that could be meaningfully compared across different arthropod orders.

## Geographic coverage

Currently, the dataset is restricted to species that are native to northwest Europe, which includes Belgium, Luxembourg, the Netherlands, northern France, United Kingdom and western Germany. Introduced species were mostly avoided, but there are some records of southern European species that have been established for a long time (e.g. *Pholcus phalangioides, Porcellionides pruinosus*).

## Taxonomic coverage

The dataset currently covers 4874 arthropod species from 10 distinct orders (Figure 2). The dataset is restricted to species with a terrestrial adult stage, which means that groups with aquatic larvae (such as Odonata) could still be incorporated into this dataset. Both the orders and species included were selected based on the availability of relevant trait data. While we aimed to be as inclusive as possible, the representation of orders and families within the dataset varies. Smaller orders like Isopoda (9 families), Odonata (9 families), Orthoptera (6 families) and Opiliones (5 families) are relatively well-covered and nearly complete. In contrast, for larger orders such as Araneae (31 families), Hemiptera (32 families) and Hymenoptera (29 families), the dataset includes about 50% of the species. Coleoptera (75 families) and Diptera (6 families) are relatively less-well represented due to a high number of species and very limited availability of trait data. Within Diptera, most data are limited to the families Syrphidae and Dolichopodidae. Lepidoptera (25 families) also appear rather incomplete when considering all species within the sampled region. However, the dataset focuses on macrolepidoptera, which are well-covered compared to the total number of species in that group.

**Figure 2:**
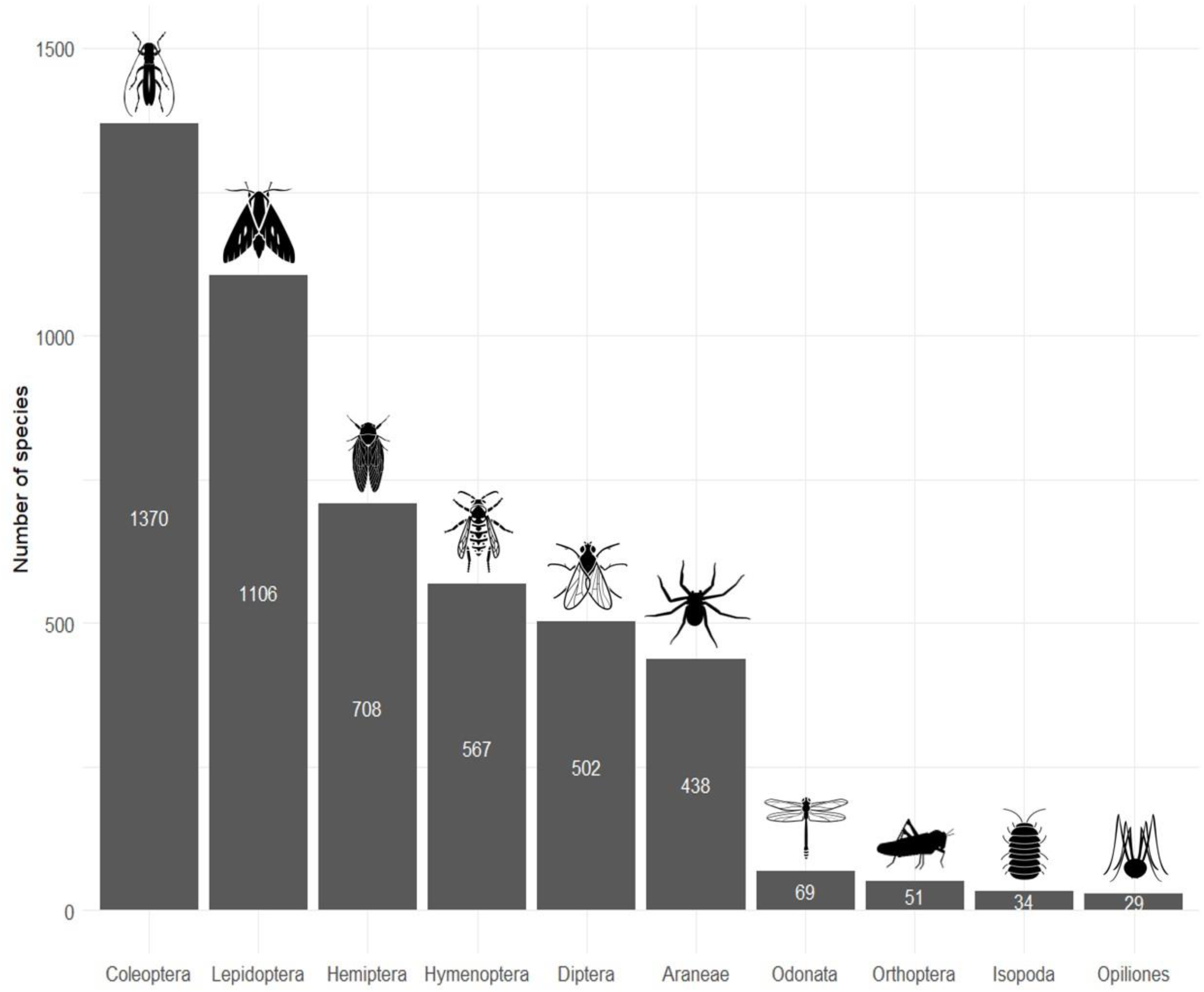
Overview of the arthropod orders that were included in the dataset. The white numbers represent the total number of species included for each order. All arthropod icons originate from Noun Project (CC BY-NC-ND 2.0). From left to right: Beetle by Rachel Siao, Moth by parkjisun, Cicada by Alejandro Capellan, wasp by parkjisun, fly by Hermine Blanquart, Spider by Matthew Davis, Dragonfly by Hermine Blanquart, Cricket by Ed Harrison, isopod by Pham Thanh Lôc and Spider by Miroslava.

No data have been included for taxa with lower taxonomic resolution than species level. For each species, other taxonomic ranks including kingdom, phylum, order, suborder and family were added to the dataset to provide fully detailed information on taxonomic position.

## Traits coverage

The dataset comprises 28 traits describing ecological and life history traits of the assessed arthropod species (Table 1). These traits were selected based on their significance in predictive modelling concerning the future effects of global change on arthropods (see above for more details).

**Table 1:**
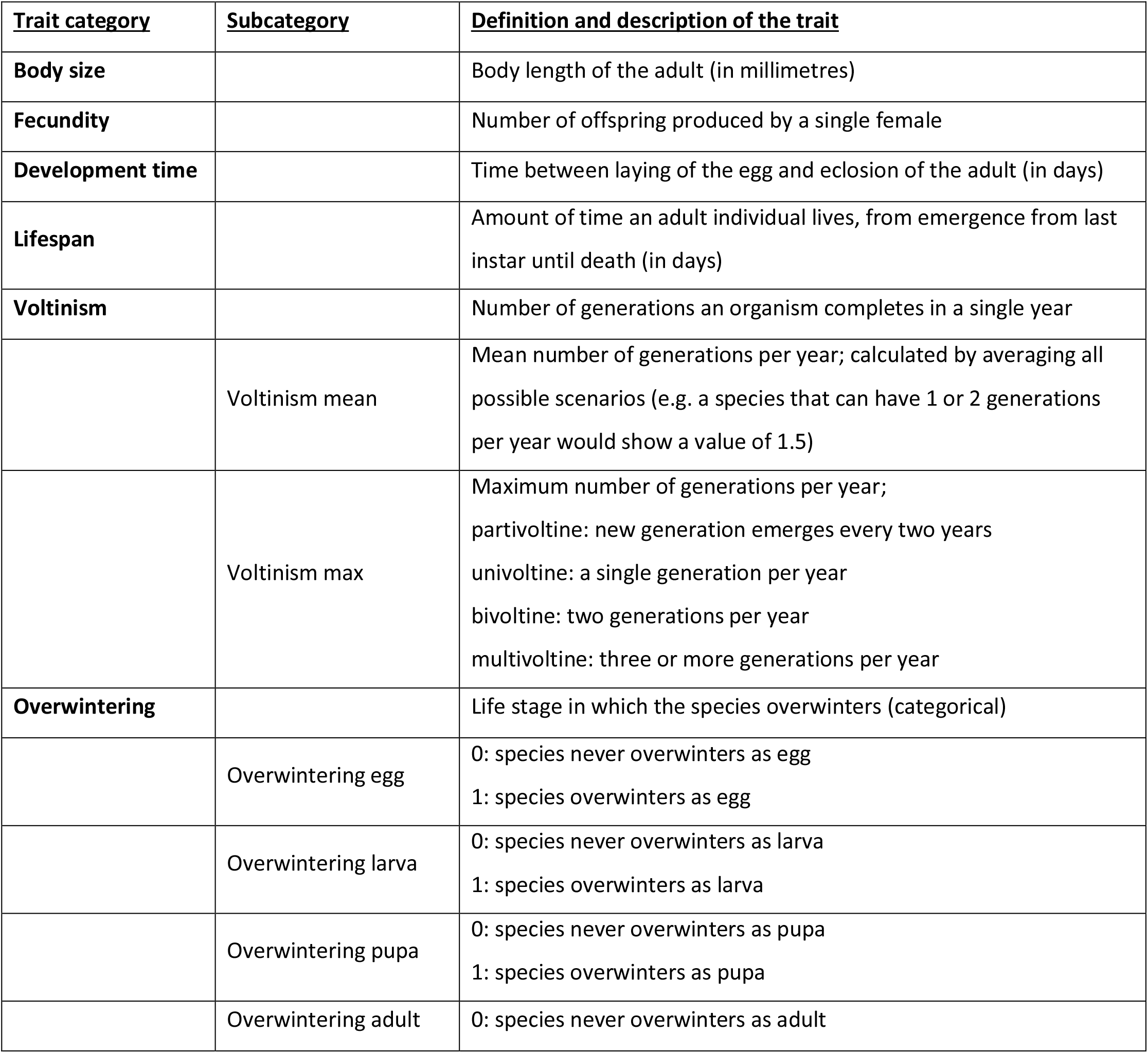
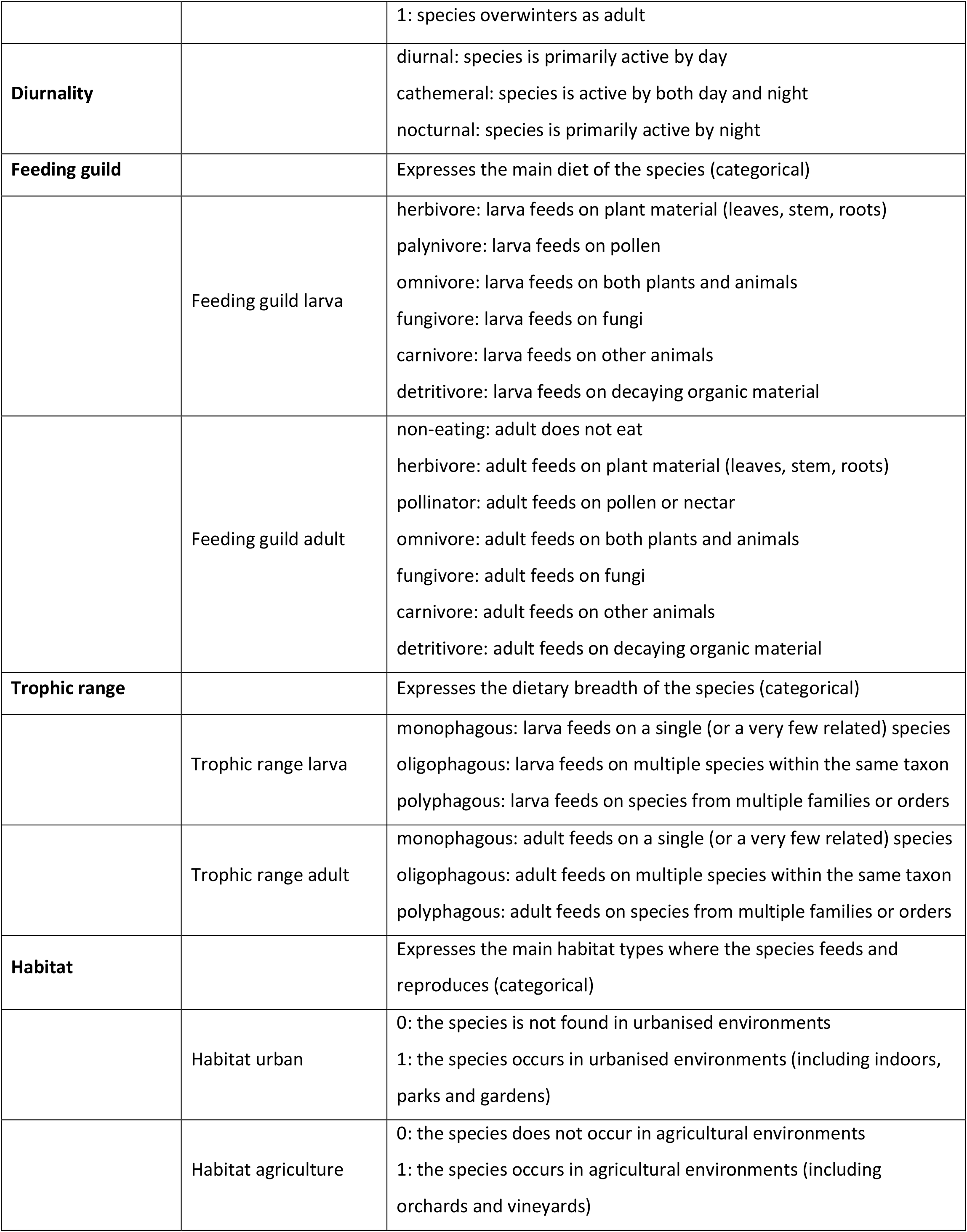
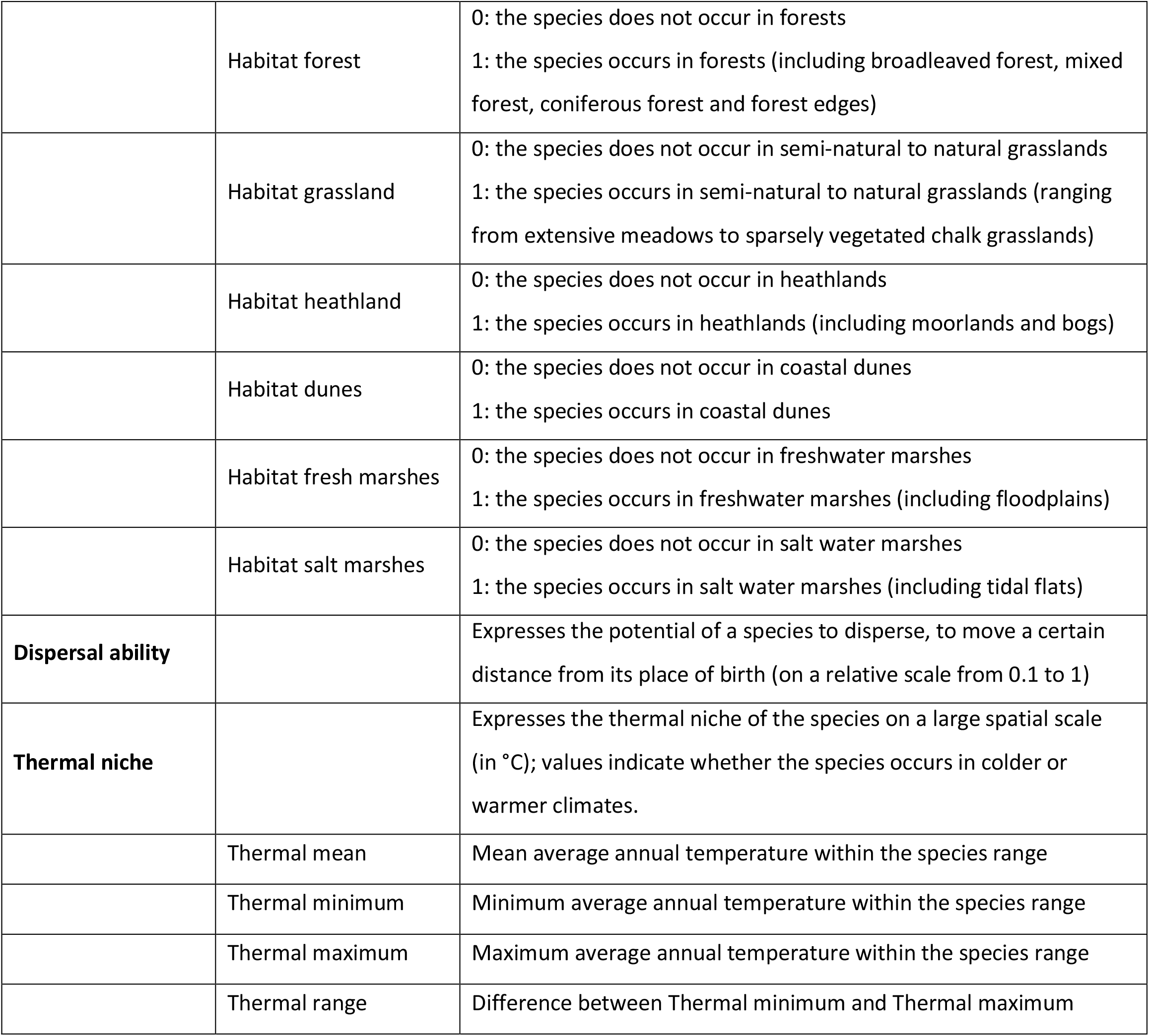
Definition of the 28 traits included in the dataset, based on (Moretti et al., 2017). In several instances, these traits constitute subcategories of broader traits. In those cases, only the subcategories have dedicated columns in the dataset.

### Data coverage of traits

The amount of available data differs substantially between traits (Figure 3). Data on body size, habitat and distribution (used for calculating thermal niche) were readily available for a wide range of species. However, data on demographic traits like development time, fecundity and lifespan were much more difficult to obtain. Most sources assessing demographic traits tend to focus on a few closely related species, causing a noticeable lack of data availability. It must be noted that even though it seems that dispersal ability has a lot of entries in the dataset, these datapoints are mostly based on proxies or expert judgement. Data on actual dispersal distances of arthropods is extremely rare, mainly due to the inherent challenges that come with tracking these small organisms (Batsleer et al., 2020).

**Figure 3:**
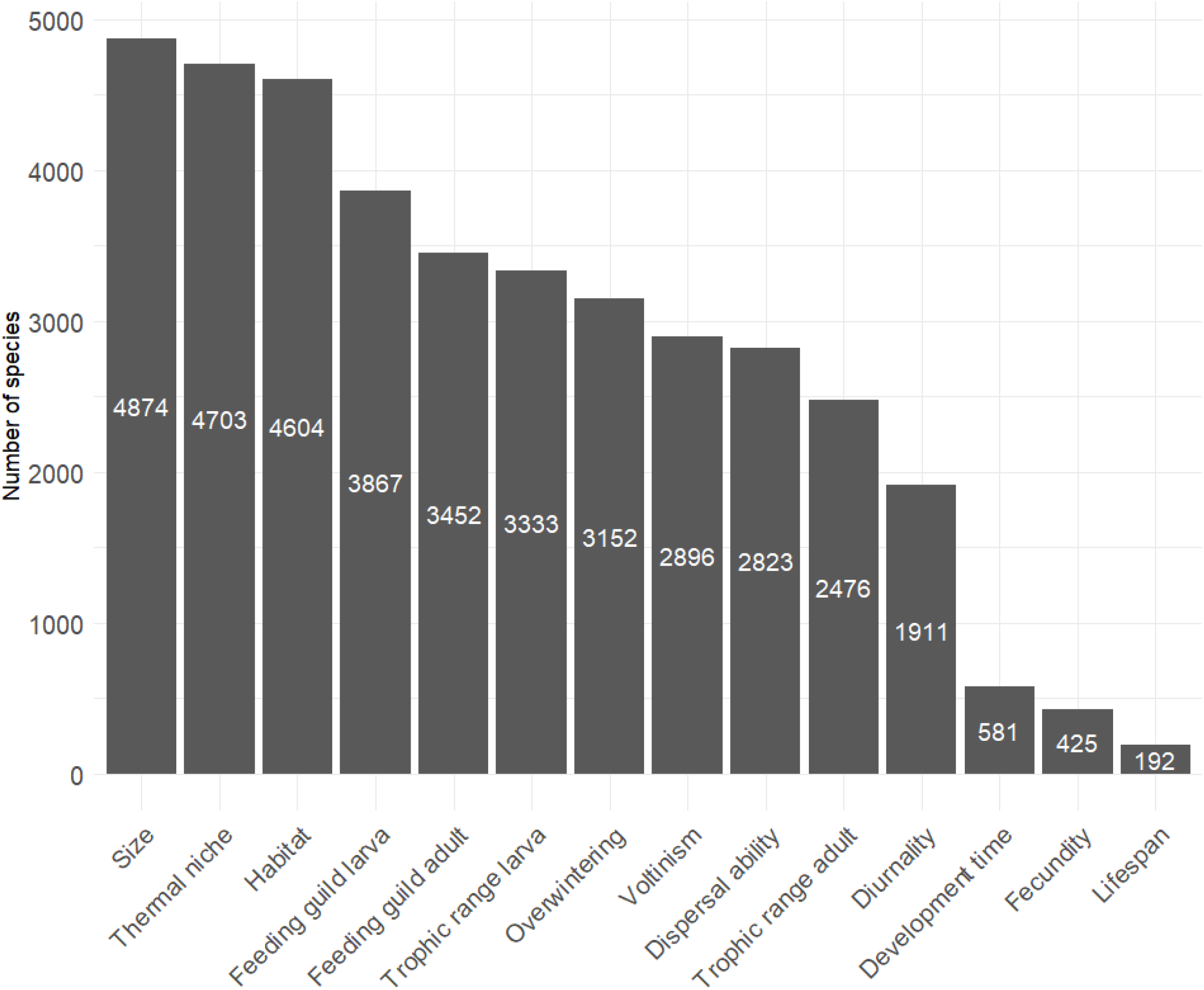
Overview of the number of species for which data on a given trait is available. The trait categories with subcategories (Habitat, Thermal niche, Overwintering & Voltinism; see table 1) have the same number of species for each subcategory for which trait data is available.

## Data resources

The final dataset was published on GBIF: https://doi.org/10.15468/75g5z9, raw datasets and code will be available from Zenodo: 10.5281/zenodo.13379714.

## Additional information

### Limitations

While we aimed to be as complete as possible, this dataset has some important limitations. First, we did not account for intraspecific variation in traits, opting instead to use averaged values for each trait. Similarly, we did not consider potential variation in average trait values across a species’ range. This decision was made because many sources report only average values, making data on trait variation difficult to obtain. Furthermore, we acknowledge there are significant limitations to our calculation of thermal niche. These data should not be used as precise thermal limits but rather as an indication whether the species is more adapted to warmer or cooler climates. While data on microclimates or physiological thermal limits would offer a more accurate estimate of thermal tolerance, they are much harder to obtain and less relevant for analysing patterns at large spatial scales and across many taxa. It is also important to note that this method calculates realised thermal niche, meaning that species might have broader thermal niches than observed but can be constricted in their distribution range by, for instance, species interactions or fragmentation (Sax et al., 2013). The accuracy of these calculations can be somewhat compromised for species with few records in GBIF. Nevertheless, we found that the values generally align well for some groups with climate niches reported in previous studies (see Supplement 2). Finally, while we present an extensive dataset covering more than 4000 species, we acknowledge that this represents only a small fraction of the total arthropod diversity in northwestern Europe. This limitation is largely due to the lack of available data for many arthropod groups, as research tends to be biased toward large and more charismatic species (Cardoso, 2012).

## Acknowledgements

The authors would like to thank the following people for providing and/or recommending literature: Diana Bowler, Wouter Dekoninck, Frederik Hendrickx, Thomas Merckx, Thomas Parmentier, Arno Thomaes, Elias Van Den Broeck, Luc Vanhercke and Pieter Vantieghem. Special thanks to Uche Osajie, who tried out the method for calculating thermal niches in QGIS and contrasted the results with available literature data.

## Author contributions

Garben Logghe, Femke Batsleer, Dirk Maes and Dries Bonte devised the content and structure of the dataset. Garben Logghe collected and synthesised the data. Tristan Permentier and Garben Logghe developed the method for calculating thermal niches. Dimitri Brosens and Stijn Cooleman coordinated the data publishing process. The following authors provided personal data and/or expert judgement for specific taxa: Matty Berg (Isopoda), Pallieter De Smedt (Isopoda), Dirk Maes (Lepidoptera), Jonas Hagge (xylophagous Coleoptera), Jorg Lambrechts (Orthoptera), Marc Pollet (Dolichopodidae) and Fons Verheyde (Hymenoptera). Garben Logghe wrote the first draft of the manuscript. All authors contributed to a first revision of the manuscript.

