## Supplementary material for "An in-depth dataset of northwestern European arthropod life histories and ecological traits"

#### S.1: list of sources consulted for data collection

Alexander, R. D. (1968). Life cycle origins, speciation, and related phenomena in crickets. *The Quarterly Review of Biology, 43*(1), 1–42.

Aukema, B., Cuppen, J. G. M., Nieser, N., & Tempelman, D. (2002). Verspreidingsatlas Nederlandse wantsen (Hemiptera: Heteroptera). Deel I: Dipsocoromorpha, Nepomorpha, Gerromorpha & Leptopodomorpha. EIS-Nederland, Leiden.

Aukema, B., Heijerman, T., & Kalkman, V. J. (2016). Veldgids wantsen deel 1. EIS-Nederland, Leiden.

Aukema, B., & Hermes, D. J. (2006). Verspreidingsatlas Nederlandse wantsen (Hemiptera: Heteroptera). Deel II: Cimicomorpha I (Tingidae, Microphysidae, Nabidae, Anthocoridae, Cimicidae & Reduviidae). EIS-Nederland, Leiden.

Aukema, B., & Hermes, D. J. (2014). Verspreidingsatlas Nederlandse wantsen (Hemiptera: Heteroptera). Deel III: Cimicomorpha II (Miridae). EIS-Nederland, Leiden.

Aukema, B., & Hermes, D. J. (2016). Verspreidingsatlas Nederlandse wantsen (Hemiptera: Heteroptera). Deel IV: Pentatomomorpha I (Aradidae, Lygaeidae, Piesmatidae, Berytidae en Pyrrhocoridae). EIS-Nederland, Leiden.

Bakewell, A. T., Davis, K. E., Freckleton, R. P., Isaac, N. J. B., & Mayhew, P. J. (2020). Comparing Life Histories across Taxonomic Groups in Multiple Dimensions: How Mammal-Like Are Insects? *The American Naturalist, 195*(1), 70–81. https://doi.org/10.1086/706195

Barbaro, L., & van Halder, I. (2009). Linking bird, carabid beetle and butterfly life history traits to habitat fragmentation in mosaic landscapes. *Ecography, 32*(2), 321–333. https://doi.org/10.1111/j.1600-0587.2008.05546.x

Baris, A. (2022). Age-Stage, Two-Sex Life Tables of the 24 Spot Ladybird [(*Subcoccinella vigintiquatuorpunctata* (Coleoptera: Coccinellidae)]. *Brazilian Archives of Biology and Technology, 65*, e22210841. https://doi.org/10.1590/1678-4324-2022210841

Bartonova, A., Benes, J., Fric, Z. F., Chobot, K., & Konvicka, M. (2016). How universal are reserve design rules? A test using butterflies and their life history traits. *Ecography, 39*(5), 456–464. https://doi.org/10.1111/ecog.01642

Biedermann, R., & Niedringhaus, R. (2009). The plant- and leafhoppers of Germany: Identification keys for all species. WABV, Germany.

Bink, F. A. (1992). Ecologische Atlas van de Dagvlinders van Noordwest-Europa. Schuyt & Co Uitgevers en Importeurs bv, Haarlem.

Boeraeve, P., Arijs, G., Segers, S., & Smedt, P. D. (2021). Habitat and seasonal activity patterns of the terrestrial arthropods. *Belgian Journal of Entomology, 116*, 1-95.

Boyes, C., & Fox, M. (2020). Bees Wasps & Ants Recording Society. https://bwars.com/

Branquart, E., & Hemptinne, J.-L. (2000). Development of ovaries, allometry of reproductive traits and fecundity of *Episyrphus balteatus* (Diptera: Syrphidae). *European Journal of Entomology, 97*(2), 165–170. https://doi.org/10.14411/eje.2000.031

British Arachnological Society. (2024). Spider and Harvestman Recording Scheme website. https://srs.britishspiders.org.uk/portal.php

Bruggink, J. (2021). *Interspecific and intraspecific variation in reproduction of terrestrial isopods.* [Master thesis] Vrije Universiteit Amsterdam.

Buse, J., Šlachta, M., Sladecek, F. X. J., & Carpaneto, G. M. (2018). Summary of the morphological and ecological traits of Central European dung beetles: Traits of Central European dung beetles. *Entomological Science, 21*(3), 315–323. https://doi.org/10.1111/ens.12313

Cook, P. M., Tordoff, G. M., Davis, A. M., Parsons, M. S., Dennis, E. B., Fox, R., Botham, M. S., & Bourn, N. A. D. (2021). Traits data for the butterflies and macro-moths of Great Britain and Ireland. *Ecology, 103*(5), e3670. https://doi.org/10.1002/ecy.3670

Cooke, J. A. L. (1965). Spider genus Dysdera (Araneae, Dysderidae). *Nature, 205*, 1027–1028.

Danks, H. V. (1968). *Bionomics of some stem-nesting aculeate Hymenoptera*. [PhD thesis] University of London.

De Smedt, P., Boeraeve, P., Arijs, G., & Segers, S. (2020). De landpissebedden van België (Isopoda: Oniscidea). Spinicornis.

De Vlinderstichting. (2024). De Vlinderstichting. Retrieved January 17, 2024, from https://www.vlinderstichting.nl/

Dekoninck, W., Vankerkhoven, F., & Maelfait, J.-P. (2003). Verspreidingsatlas en voorlopige Rode Lijst van de mieren van Vlaanderen. Institute for Nature and Forest (INBO).

Dziock, F., Gerisch, M., Siegert, M., Hering, I., Scholz, M., & Ernst, R. (2011). Reproducing or dispersing? Using trait based habitat templet models to analyse Orthoptera response to flooding and land use. *Agriculture, Ecosystems & Environment, 145*(1), 85–94. https://doi.org/10.1016/j.agee.2011.07.015

Ellis, W. (2020). Plantparasieten van Europa: Bladmineerders, gallen en schimmels. https://bladmineerders.nl/?lang=nl

Franzén, M., & Nilsson, S. G. (2012). Climate-dependent dispersal rates in metapopulations of burnet moths. *Journal of Insect Conservation, 16*(6), 941–947. https://doi.org/10.1007/s10841-012-9481-4

García-Barros, E. (2000). Body size, egg size, and their interspecific relationships with ecological and life history traits in butterflies (Lepidoptera: Papilionoidea, Hesperioidea). *Biological Journal of the Linnean Society, 70*(2), 251–284. https://doi.org/10.1111/j.1095-8312.2000.tb00210.x

Geden, C. J. (1984). Population dynamics, spatial distribution, dispersal behavior and life history of the predaceous histerid, *Carcinops pumilio* (Erichson), with observations of other members of the poultry manure arthropod community. [PhD thesis] University of Massachusetts Amherst.

Gittings, T., & Giller, P. S. (1997). Life history traits and resource utilisation in an assemblage of north temperate *Aphodius* dung beetles (Coleoptera: Scarabaeidae). *Ecography, 20*(1), 55–66. https://doi.org/10.1111/j.1600-0587.1997.tb00347.x

Gossner, M. M., Simons, N. K., Achtziger, R., Blick, T., Dorow, W. H. O., Dziock, F., Köhler, F., Rabitsch, W., & Weisser, W. W. (2015). A summary of eight traits of Coleoptera, Hemiptera, Orthoptera and Araneae, occurring in grasslands in Germany. *Scientific Data, 2*, 150013. https://doi.org/10.1038/sdata.2015.13

Haack, R. A., Keena, M. A., & Eyre, D. (2017). Life History and Population Dynamics of Cerambycids. In: *Wang (2017). Cerambycidae of the World: Biology and Pest Management.* (pp. 71-103) CRC Press, Boca Raton, Florida.

Haes, E. C. M., & Harding, P. T. (1997). Atlas of grasshoppers, crickets, and allied insects in Britain and Ireland. Stationery Office, London.

Harabis, F., & Hronkova, J. (2020). European database of the life history, morphological and habitat characteristics of dragonflies (Odonata). *European Journal of Entomology, 117*, 302–308. https://doi.org/10.14411/eje.2020.035

Hodek, I., & Honěk, A. (1996). Ecology of Coccinellidae. Springer Netherlands, Dordrecht. https://doi.org/10.1007/978-94-017-1349-8

Jones, H. B. C., Lim, K. S., Bell, J. R., Hill, J. K., & Chapman, J. W. (2016). Quantifying interspecific variation in dispersal ability of noctuid moths using an advanced tethered flight technique. *Ecology and Evolution, 6*(1), 181–190. https://doi.org/10.1002/ece3.1861

Kalkman, V. J., Aukema, B., & Heijerman, T. (2015). Soortzoeker Netwantsen van Nederland. Naturalis Biodiversity Center & EIS Kenniscentrum Insecten, Leiden.

Kleukers, R. (2017). De sprinkhanen van Nederland en België. Jeugdbondsuitgeverij, ‘s Gravenland.

Kleukers, R., Van Nieukerken, E., Ode, B., Willemse, L., & Van Wingerden, W. (2004). De sprinkhanen en krekels van Nederland. Naturalis, EIS-Nederland, KNNV, Leiden.

Komonen, A., Grapputo, A., Kaitala, V., Kotiaho, J. S., & Päivinen, J. (2004). The role of niche breadth, resource availability and range position on the life history of butterflies. *Oikos, 105*(1), 41–54. https://doi.org/10.1111/j.0030-1299.2004.12958.x

Kuhlmann, U. (1995). Biology of *Triathria setipennis* (Fallén) (Diptera: Tachinidae), a native parasitoid of the European earwig *Forficula auricularia* L. (Dermaptera: Forficulidae), in Europe. *The Canadian Entomologist, 127*, 507–517.

Majerus, M. E. N. (1994). Ladybirds (Vol. 81). HarperCollins, Madison, Winconsin.

Merckx, T., Feber, R. E., Mclaughlan, C., Bourn, N. A. D., Parsons, M. S., Townsend, M. C., Riordan, P., & Macdonald, D. W. (2010). Shelter benefits less mobile moth species: The field-scale effect of hedgerow trees. *Agriculture, Ecosystems & Environment, 138*(3–4), 147–151. https://doi.org/10.1016/j.agee.2010.04.010

Merckx, T., Feber, R. E., Parsons, M. S., Bourn, N. A. D., Townsend, M. C., Riordan, P., & Macdonald, D. W. (2010). Habitat preference and mobility of *Polia bombycina*: Are non-tailored agri-environment schemes any good for a rare and localised species? *Journal of Insect Conservation*, *14*(5), 499–510. https://doi.org/10.1007/s10841-010-9279-1

Middleton-Welling, J., Dapporto, L., García-Barros, E., Wiemers, M., Nowicki, P., Plazio, E., Bonelli, S., Zaccagno, M., Šašić, M., Liparova, J., Schweiger, O., Harpke, A., Musche, M., Settele, J., Schmucki, R.,

& Shreeve, T. (2020). A new comprehensive trait database of European and Maghreb butterflies, Papilionoidea. *Scientific Data, 7*(1), 351. https://doi.org/10.1038/s41597-020-00697-7

Mo, J., Baker, G., Keller, M., & Roush, R. (2003). Local Dispersal of the Diamondback Moth (*Plutella xylostella* (L.)) (Lepidoptera: Plutellidae). *Environmental Entomology, 32*(1), 71–79. https://doi.org/10.1603/0046-225X-32.1.71

Muilwijk, J., Felix, R., Dekoninck, W., & Bleich, O. (2015). De loopkevers van Nederland en België (Carabidae) (Vol. 9). EIS, Leiden.

Nentwig, W., Blick, T., Bosmans, R., Gloor, D., Hänggi, A., & Kropf, C. (2024). Spiders of Europe. Version 09.2024. Online at https://www.araneae.nmbe.ch, accessed on 03/09/2024. https://doi.org/10.24436/1

Nickel, H. (2003). The leafhoppers and planthoppers of Germany (Hemiptera, Auchenorrhyncha): Patterns and strategies in a highly diverse group of phytophagous insects. Pensoft, Sofia.

Peeters, T. M. J., Nieuwenhuijsen, H., Smit, J., van der Meer, F., Kwak, M., Loonstra, A. J., de Rond, J., Roos, M., & Reemer, M. (2012). De Nederlandse bijen (Hymenoptera: Apidae s.l.). Naturalis, EIS-Nederland, KNNV, Leiden.

Peeters, T. M. J., van Achterberg, C., Heitmans, W. R. B., Klein, W. F., Lefeber, V., van Loon, A. J., Mabelis, A. A., Nieuwenhuijsen, H., Reemer, M., de Rond, J., Smit, J., & Velthuis, H. H. W. (2004). De wespen en mieren van Nederland (Hymenoptera: Aculeata). Naturalis, EIS-Nederland, KNNV, Leiden.

Pekár, S. (2000). Webs, diet, and fecundity of *Theridion impressum* (Araneae: Theridiidae). *European Journal of Entomology, 97*(1), 47–50.

Pekár, S., Coddington, J. A., & Blackledge, T. A. (2012). Evolution of stenophagy in spiders (Araneae): evidence based on the comparative analysis of spiders diets. *Evolution, 66*(3), 776–806. https://doi.org/10.1111/j.1558-5646.2011.01471.x

Pekár, S., & Jarab, M. (2011). Life history constraints in inaccurate Batesian myrmecomorphic spiders (Araneae: Corinnidae, Gnaphosidae). *European Journal of Entomology, 108*(2), 255–260. https://doi.org/10.14411/eje.2011.034

Pekár, S., Wolff, J. O., Černecká, Ľ., Birkhofer, K., Mammola, S., Lowe, E. C., Fukushima, C. S., Herberstein, M. E., Kučera, A., Buzatto, B. A., Djoudi, E. A., Domenech, M., Enciso, A. V., Piñanez Espejo, Y. M. G., Febles, S., García, L. F., Gonçalves-Souza, T., Isaia, M., Lafage, D., … Cardoso, P. (2021). The World Spider Trait database: A centralized global open repository for curated data on spider traits. Database, 2021, baab064. https://doi.org/10.1093/database/baab064

Pitts, K. M., & Wall, R. (2004). Adult mortality and oviposition rates in field and captive populations of the blowfly *Lucilia sericata*. *Ecological Entomology, 29*(6), 727–734. https://doi.org/10.1111/j.0307-6946.2004.00653.x

Pont, & Meier, R. (2002). The Sepsidae (Diptera) of Europe. BRILL, Leiden. https://doi.org/10.1163/9789047401391

Potocký, P., Bartoňová, A., Beneš, J., Zapletal, M., & Konvička, M. (2018). Life history traits of Central European moths: Gradients of variation and their association with rarity and threats. *Insect Conservation and Diversity*, *11*(5), 493–505. https://doi.org/10.1111/icad.12291

Rahman, M. A., Sarker, S., Ham, E., Lee, J.-S., & Lim, U. T. (2020). Development and Fecundity of *Orius minutus* (Hemiptera: Anthocoridae) and *O. laevigatus* reared on *Tetranychus urticae* (Acari: Tetranychidae). *Journal of Economic Entomology, 113*(4), 1735–1740. https://doi.org/10.1093/jee/toaa078

Reemer, M., Renema, W., van Steenis, W., Zeegers, T., Barendregt, A., Smit, J. T., van Veen, M. P., van Steenis, J., & van der Leij, L. J. J. M. (2009). De Nederlandse zweefvliegen (Diptera: Syrphidae). Naturalis, EIS-Nederland, KNNV, Leiden.

Rheinheimer, J., & Hassler, M. (2010). Die Rüsselkäfer Baden-Württembergs (Vol. 99). Landesantalt für Umwelt Baden-Württemberg, Karlsruhe.

Ribera, I., Dolédec, S., Downie, I. S., & Foster, G. N. (2001). Effect of land disturbance and stress on species traits of ground beetle assemblages. *Ecology, 82*(4), 1112–1129. https://doi.org/10.1890/0012-9658(2001)082[1112:EOLDAS]2.0.CO;2

Rigal, F., Cardoso, P., Lobo, J. M., Triantis, K. A., Whittaker, R. J., Amorim, I. R., & Borges, P. A. V. (2018). Functional traits of indigenous and exotic ground-dwelling arthropods show contrasting responses to land-use change in an oceanic island, Terceira, Azores. *Diversity and Distributions, 24*(1), 36–47. https://doi.org/10.1111/ddi.12655

Rouault, G., Cantini, R., Battisti, A., & Roques, A. (2005). Geographic distribution and ecology of two species of *Orsillus* (Hemiptera: Lygaeidae) associated with cones of native and introduced Cupressaceae in Europe and the Mediterranean Basin. *The Canadian Entomologist, 137*(4), 450–470. https://doi.org/10.4039/n04-044

Ruhland, F., Pétillon, J., & Trabalon, M. (2016). Physiological costs during the first maternal care in the wolf spider *Pardosa saltans* (Araneae, Lycosidae). *Journal of Insect Physiology, 95*, 42–50. https://doi.org/10.1016/j.jinsphys.2016.09.007

Schweiger, A. H., & Beierkuhnlein, C. (2016). Size dependency in colour patterns of Western Palearctic carabids. *Ecography, 39*(9), 846–857. https://doi.org/10.1111/ecog.01570

Segers, S. (2015). Velddeterminatietabel voor de lieveheersbeestjes van West-Europa (Chilocorinae, Coccinellinae, Epilachninae & Coccidulinae). Jeugdbond voor Natuur en Milieu, Gent.

Slade, E. M., Merckx, T., Riutta, T., Bebber, D. P., Redhead, D., Riordan, P., & Macdonald, D. W. (2013). Life history traits and landscape characteristics predict macro-moth responses to forest fragmentation. *Ecology, 94*(7), 1519–1530. https://doi.org/10.1890/12-1366.1

Spitzer, K., Rejmánek, M., & Soldán, T. (1984). The fecundity and long-term variability in abundance of noctuid moths (Lepidoptera, Noctuidae). *Oecologia, 62*(1), 91–93. https://doi.org/10.1007/BF00377379

Staab, M., Gossner, M. M., Simons, N. K., Achury, R., Ambarlı, D., Bae, S., Schall, P., Weisser, W. W., & Blüthgen, N. (2023). Insect decline in forests depends on species' traits and may be mitigated by management. *Communications Biology, 6*(1), 338. https://doi.org/10.1038/s42003-023-04690-9

Stevens, V. M., Whitmee, S., Le Galliard, J.-F., Clobert, J., Böhning-Gaese, K., Bonte, D., Brändle, M., Matthias Dehling, D., Hof, C., Trochet, A., & Baguette, M. (2014). A comparative analysis of dispersal syndromes in terrestrial and semi-terrestrial animals. *Ecology Letters, 17*(8), 1039–1052. https://doi.org/10.1111/ele.12303

Thiele, H.-U. (1977). Carabid Beetles in Their Environments. Springer Berlin Heidelberg. https://doi.org/10.1007/978-3-642-81154-8

Torres-Vila, L. M. (2017). Reproductive biology of the great capricorn beetle, *Cerambyx cerdo* (Coleoptera: Cerambycidae): a protected but occasionally harmful species. *Bulletin of Entomological Research, 107*(6), 799–811. https://doi.org/10.1017/S0007485317000323

Turin, H. (2000). De Nederlandse loopkevers, verspreiding en oecologie (Coleoptera: Carabidae): Vol. Nederlandse Fauna (3rd ed.). KNNV Uitgeverij & EIS-Nederland, Leiden.

UK Beetles. (2024). UK Beetles. Retrieved January 11, 2024, from https://www.ukbeetles.co.uk/

van den Broek, R., & Schulten, A. (2013). Roofvliegen van Nederland en België inclusief Denemarken en de Britse Eilanden (Diptera—Asilidae).

van Noordwijk, T., & Siepel, H. (2014). *Through arthropod eyes: Gaining mechanistic understanding of calcareous grassland diversity.* [PhD thesis ] Ghent University.

Vibert, S., Salomon, M., Scott, C., Blackburn, G. S., & Gries, G. (2017). Life history data for the funnel weavers *Eratigena agrestis* and *Eratigena atrica* (Araneae: Agelenidae) in the Pacific Northwest of North America. *The Canadian Entomologist, 149*(3), 345–356. https://doi.org/10.4039/tce.2016.73

Wagner, W. (2005). Schmetterlinge und ihre Ökologie. http://pyrgus.de/

Waloff, N. (2009). The parasitoids of the nymphal and adult stages of leafhoppers (Auchenorrhyncha: Homoptera) of acidic grassland. *Transactions of the Royal Entomological Society of London, 126*(4), 637–686. https://doi.org/10.1111/j.1365-2311.1975.tb00862.x

Wang, X., Zuo, L., Yun, Y., Chen, J., Zhang, Z., & Peng, Y. (2016). The responses of a funnel-web weaving spider, *Agelena labyrinthica*, to elevated CO 2 concentration. *Entomologia Experimentalis et Applicata, 161*(3), 213–218. https://doi.org/10.1111/eea.12511

Wijnhoven, H. (2009). De Nederlandse hooiwagens (Opiliones) (Vol. 3). EIS-Nederland, Leiden.

Winkelman, J. (2013). De Nederlandse Goudhaantjes (Chrysomelidae: Chrysomelinae) (Vol. 7). EIS-Nederland, Leiden.

Zeegers, T., & Heijerman, T. (2008). De Nederlandse boktorren (Cerambycidae) (Vol. 2). EIS-Nederland, Leiden.

### S.2: linear regressions between temperature estimates

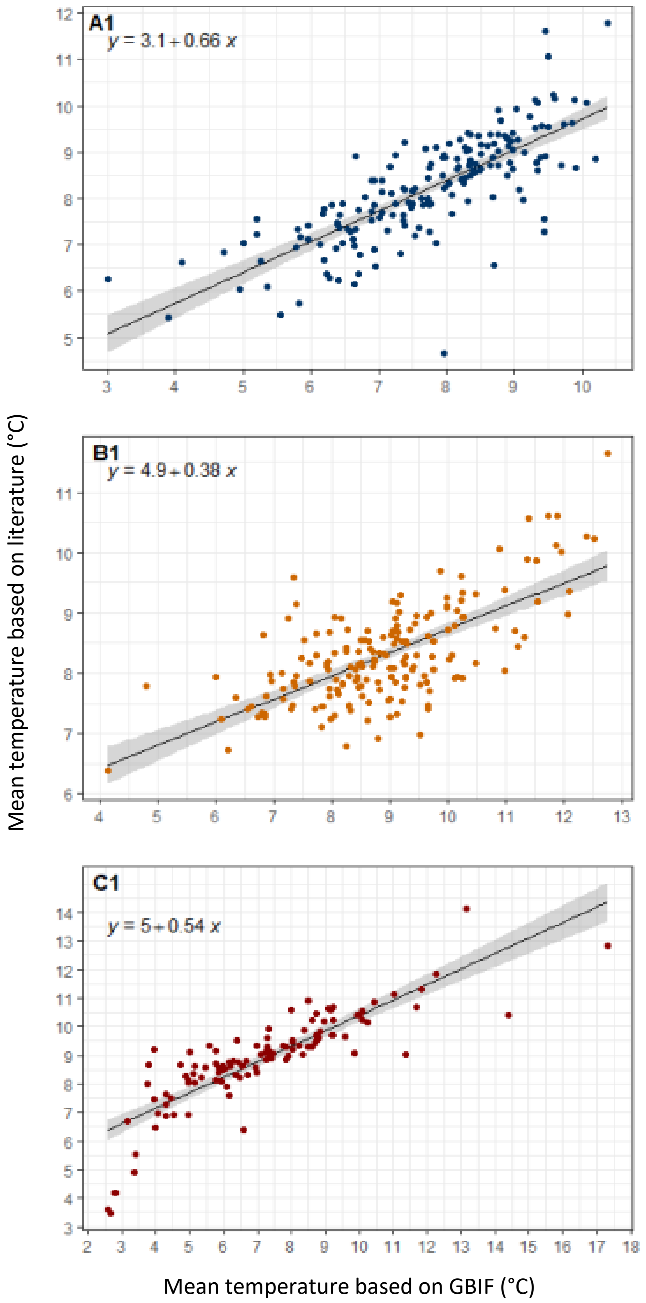


*Figure S.2: Linear regressions between temperature estimates from literature and our calculations based on GBIF distribution data. Three different arthropod groups were assessed based on available literature data: (A) Carabidae, (B) Araneae and (C) Rhopalocera. Literature values were derived from Schweiger et al. (2014) and Bowler et al. (2017). The figure is adapted from the bachelor thesis of Uche Osajie (2021).*
